# Pathways controlling neurotoxicity and proteostasis in mitochondrial complex I deficiency

**DOI:** 10.1101/2024.01.08.574634

**Authors:** Vanitha Nithianadam, Souvarish Sarkar, Mel B. Feany

## Abstract

Neuromuscular disorders caused by dysfunction of the mitochondrial respiratory chain are common, severe and untreatable. We recovered a number of mitochondrial genes, including electron transport chain components, in a large forward genetic screen for mutations causing age-related neurodegeneration in the context of proteostasis dysfunction. We created a model of complex I deficiency in the *Drosophila* retina to probe the role of protein degradation abnormalities in mitochondrial encephalomyopathies. Using our genetic model, we found that complex I deficiency regulates both the ubiquitin/proteasome and autophagy/lysosome arms of the proteostasis machinery. We further performed an in vivo kinome screen to uncover new and potentially druggable mechanisms contributing to complex I related neurodegeneration and proteostasis failure. Reduction of RIOK kinases and the innate immune signaling kinase pelle prevented neurodegeneration in complex I deficiency animals. Genetically targeting oxidative stress, but not RIOK1 or pelle knockdown, normalized proteostasis markers. Our findings outline distinct pathways controlling neurodegeneration and protein degradation in complex I deficiency and introduce an experimentally facile model in which to study these debilitating and currently treatment-refractory disorders.

## Introduction

Mitochondrial electron transport disorders are common human genetic diseases, with a collective estimated prevalence of approximately 1 in 5,000 (1). Mutations linked to human disease have been described in oxidative phosphorylation enzymes in complexes I, II, III, cytochrome c oxidase (complex IV), and ATP synthase (complex V), as well as in assembly factors for each of the enzyme complexes (2). Clinical phenotypes resulting from electron transport chain deficiency are protean and include early onset, severe necrotizing leukoencephalitis (Leigh’s syndrome), parkinsonism, late adult-onset retinal degeneration (Leber’s hereditary optic neuropathy) and myopathies (3). Deficiencies in complex I are the most common cause of single mitochondrial enzyme deficiency, perhaps reflecting the large size of the complex I enzyme (4). The complex is made up of 44 subunits, seven encoded by mitochondrial genes and 37 by nuclear genes. Disease-causing mutations have been described in 7 mitochondrial and 26 nuclear genes. Mutations in 9 of the genes encoding the 15 putative complex I assembly factors have also been implicated in disease (5).

The overall function of the respiratory chain is to transduce energy from cellular metabolism into ATP. During oxidative phosphorylation energy from electron transport is converted by complexes I, III and IV into an electrochemical proton gradient, which is converted into ATP by complex V (ATPase). Complex II (succinate dehydrogenase) contributes to electron transfer by reducing ubiquinone. Defective energy production has been implicated in the pathogenesis of complex I deficiency and other mitochondrial disorders, consistent with the tendency for preferential involvement of organs with high energy demands, including brain and cardiac and skeletal muscle (2). Disruption in mitochondrial respiration also leads to excess production of reactive oxygen species. Unfortunately, despite these mechanism-linked targets for therapeutic intervention effective therapies for patients with complex I deficiency and other mitochondrial diseases remain an important unmet clinical need (6).

To facilitate new approaches to understanding the pathogenesis of electron transport chain deficiency, identify novel therapeutic targets and allow drug screening in vivo, a number of *Drosophila* models have been developed of complex I deficiency. Global reduction of the mitochondrially encoded core ND2 subunit (7) or the nuclear encoded NDUFS8 core subunit (ND-23 in *Drosophila*) leads to reduced lifespan and neurodegeneration (8). The ND2 model has additionally been used to suggest a beneficial effect of rapamycin treatment (9), consistent with positive effects of rapamycin in murine models of complex I deficiency (10) and mitochondrial depletion (11). Studies with NDUFS8 mutants implicated interactions with the maternally inherited mitochondrial genome as modifiers of disease severity (8).

We have previously shown that global knockdown of the core NDUFV1 subunit (ND-75) in the brain reduces lifespan, impairs locomotor function and causes neurodegeneration (12). Interestingly, glial complex I knockdown is critical for neurodegeneration in our previously described model. Similar findings of decreased longevity and impaired locomotion have been reported following neuronal and glial knockdown of the core NDUFS8 subunit (ND-23) (13). NDUFS8 knockdown was accompanied by accumulation of lipid droplets in glia, as was previously observed in retinal pigment cells following disruption of mitochondrial function (14).

Although these efforts have led to intriguing mechanistic insights and potential therapeutic leads, we and others have observed that global, neuronal- or glial-specific knockdown of complex I subunits can lead to significant toxicity (15), hampering higher throughput approaches to understanding disease pathogenesis and identifying treatments such as genetic (16) or drug (17) screening. The current work describes flexible new retinal toxicity models of complex I deficiency and uses these models to identify novel potential therapeutic kinase targets for the disorder.

## Results

As part of our ongoing efforts to understand the molecular mechanisms underlying maintenance of neuronal viability and normal proteostasis during aging we knocked down 6,258 genes conserved from *Drosophila* to humans in neurons with transgenic RNAi using the *elav-GAL4* pan-neuronal driver (18). Animals were aged to 30 days and microscopic examination of brains was performed following formalin fixation and paraffin embedding (Fig. 1A). Histologic sections were assessed for evidence of neurodegeneration following hematoxylin and eosin staining and were also immunostained with an antibody recognizing ubiquitin to identify protein aggregates (19,20). We recovered a number of mitochondrial genes as preliminary hits causing neurodegeneration with accumulation of ubiquitin-positive aggregates, including genes encoding the cytochrome c assembly factor COX10, and the complex I subunits NDUFV1 (Fig. 1B, arrows, and C) and NDUFS7. Mutations in the mitochondrially encoded cytochrome c assembly factor COX10 cause Leigh’s disease, a mitochondrial encephalopathy (21), while NDUFV1 is the largest (75 kDa) subunit of complex I. These data suggest that mitochondrial function is critical for maintaining proteostasis during aging, and further that abnormal proteostasis might contribute to the pathogenesis of mitochondrial encephalomyopathies.

**Figure 1.**
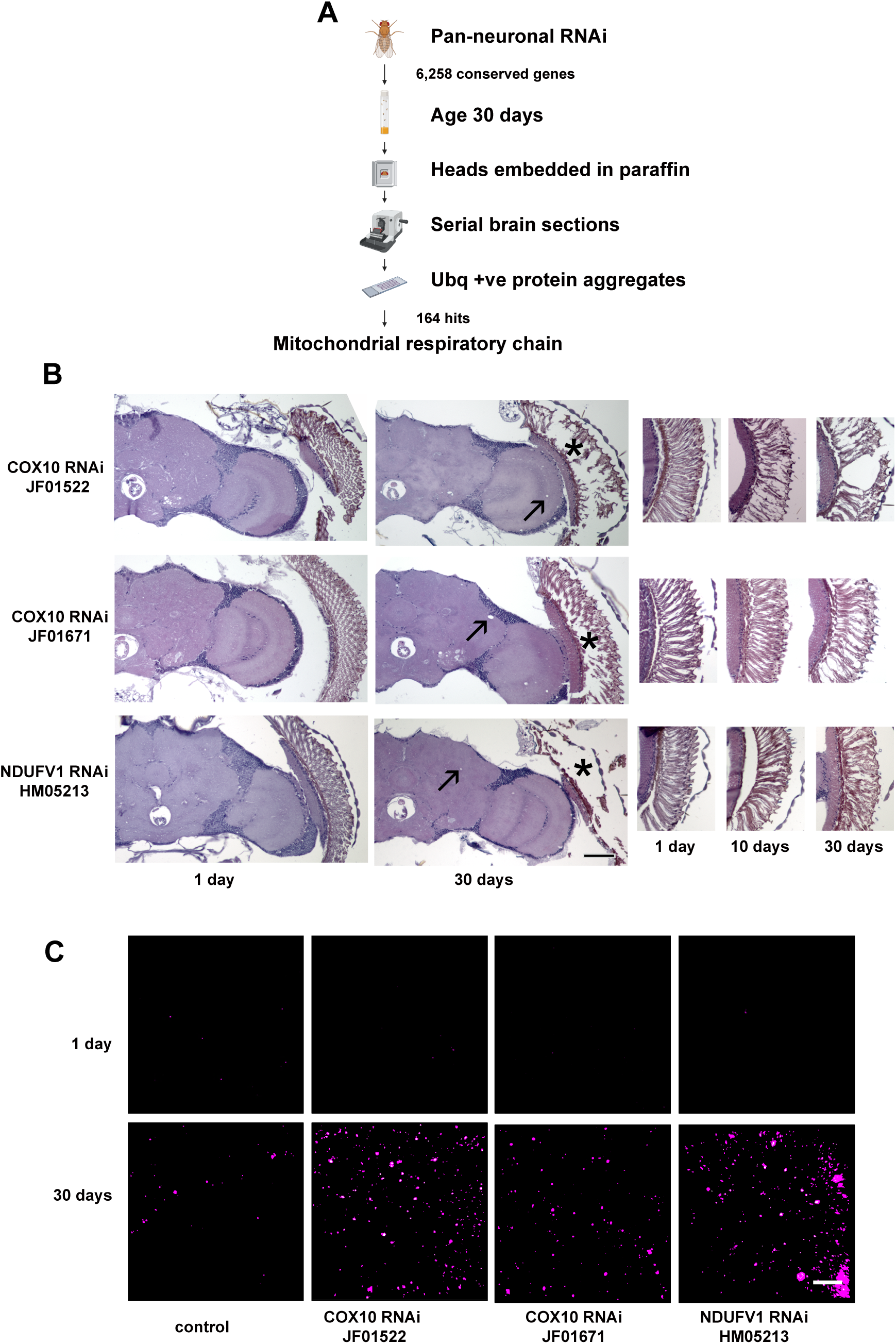
Neurodegeneration and dysproteostasis produced by mitochondrial dysfunction during aging. (A) Schematic diagram of the screening strategy designed to identify genes required for maintenance of neuronal viability and proteostasis during aging and outcome of the genome-scale neuronal transgenic RNAi knockdown screen. (B) Hematoxylin and eosin-stained sections of brains (left) with pan-neuronal knockdown of COX10 or NDUFV1 show increased vacuolization, arrows, a sign of neurodegeneration in *Drosophila*, and retinal degeneration, asterisks, with age. Cross sections of retinas (right) illustrate progressive degeneration with increased age. (C) Ubiquitin staining in the medullary neuropil in the brain of flies with knockdown of COX10 or NDUFV1 shows increased numbers of protein aggregates with age and further increases with knockdown of respiratory chain components. Scale bars are 50 µm in (A) and 5 µm in (B). Control is *elav-GAL4/+; UAS-Dcr2/+.* Flies are the indicated ages. Alt text: Schematic diagram of pan-neuronal transgenic RNAi screen and photographs showing brain and retinal degeneration and accumulation of ubiquitin-positive aggregates with knockdown of genes encoding mitochondrial respiratory chain components.

Since complex I deficiency is the most common cause of electron transport chain deficiency, we focused our subsequent efforts on NDUFV1. As previously described (15), we found that global, and widespread neuronal- and muscle-specific knockdown produced lethality or excessive toxicity with most available transgenic RNAi lines. Retinal degeneration was prominent in our pan-neuronal knockdown studies (Fig. 1B, asterisks, and Supplementary Fig. 1) and is a frequent manifestation of mitochondrial disorders in patients. We thus turned to retina-specific knockdown of NDUFV1 mediated by the *GMR (glass multimer reporter)-GAL4* driver. Retinal knockdown is also an attractive experimental approach because eyes are not required for viability in *Drosophila* and assessment of retinal phenotypes is straightforward (22,23). Consistent with our findings with pan-neuronal knockdown, when we knocked down NDUFV1 in the retina using three independent RNAi lines we observed age-dependent retinal degeneration (Supplementary Fig. 1).

We next determined if inhibition of complex I leading to retinal degeneration was accompanied by altered proteostasis in vivo by assessing the activity of the two major cellular protein degradation pathways: the ubiquitin/proteasome and autophagy/lysosome systems. To assay proteasome activity in vivo we used two fluorescent reporters. The first, GFP-CL1 is a fusion protein generated by introducing a degradation signal to GFP, which is rapidly degraded by the proteasome. Levels of the reporter have therefore been used to assess the functionality of the ubiquitin/proteasome system (24,25). We crossed the GFP-CL1 reporter flies to animals with NDUFV1 knockdown in the retina and assessed transgene levels by immunofluorescence (Fig. 2A) and western blot (Fig. 2B and C) using monoclonal antibodies directed to GFP. We observed significant elevation of GFP-CL1 in knockdown animals using both detection methods, compatible with decreased proteasome activity following complex I inhibition. To confirm our results we used a second reporter line for expression of GFP, expressing a form of GFP destabilized with the G76V point mutation (24). Again, NDUFV1 knockdown resulted in increased levels of the unstable GFP variant (Fig. 2D).

**Figure 2.**
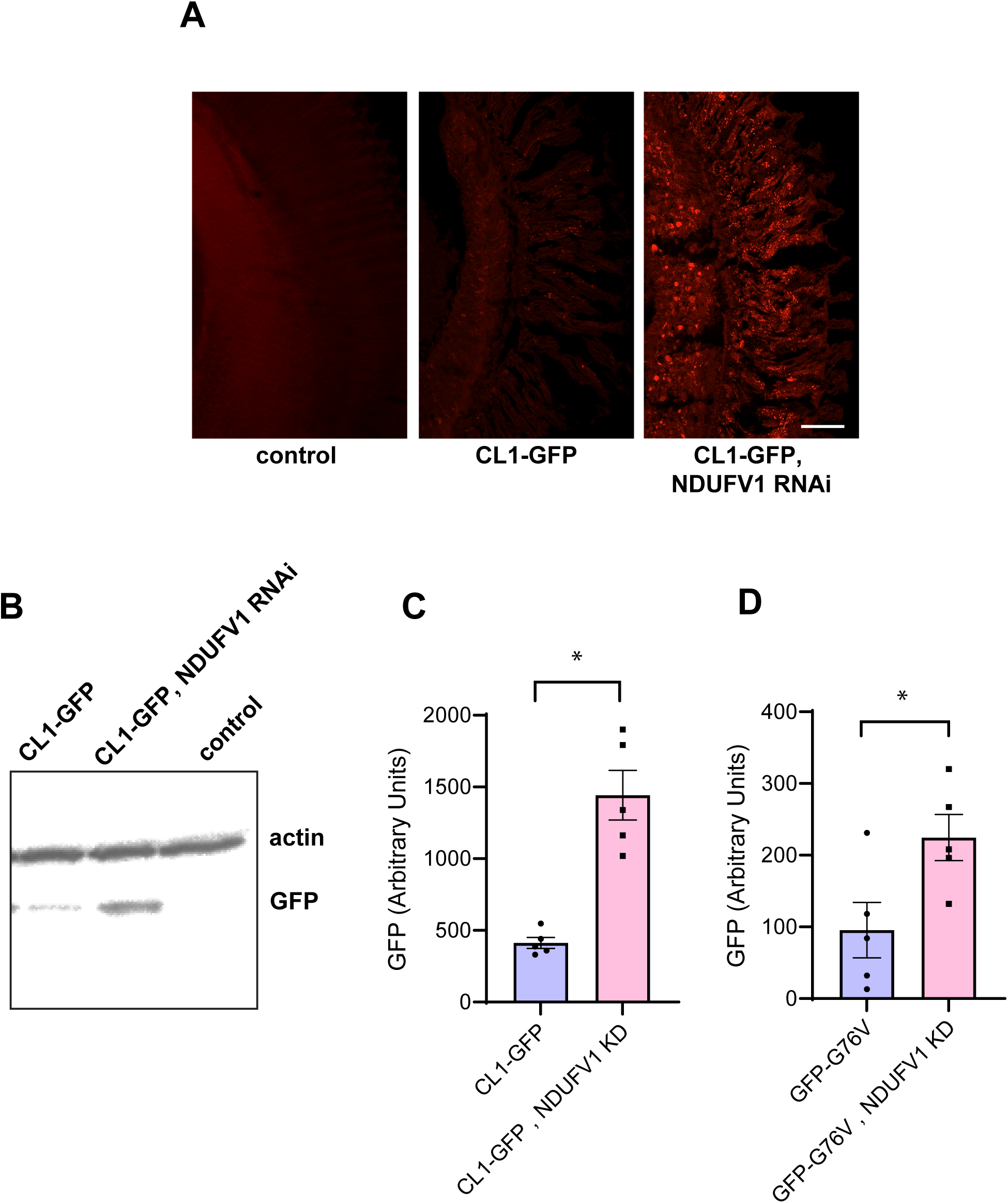
Reporters of proteasome function suggest reduced proteosome degradation following NDUFV1 knockdown. (A) Immunostaining for GFP in retinas from control flies and flies expressing the unstable CL1-GFP reporter with and without retinal knockdown of NDUFV1 shows increased levels of unstable GFP following complex I inhibition. (B,C) Western blot (B) with quantification (C) shows increased levels of the CL1-GFP reporter with retinal NDUFV1 knockdown. The blot is reprobed with an antibody to actin to illustrate equivalent protein loading. (D) Western blot with quantification shows increased levels of the GFP-G76V reporter with NDUFV1 knockdown. n = 5 per genotype. Data are represented as mean ± SEM. * p<0.01, t-test. Control is *GMR-GAL4/+.* Flies are 10 days old. Alt text: Photographs, western blot and bar graphs with quantification showing increased stability of unstable GFP variants following NDUFV1 knockdown.

To determine if the other major cellular protein degradation pathway, the autophagy/lysosome system, was affected by complex I inhibition we first assessed levels of the autophagy-related gene 8a (Atg8a) protein, the fly homolog of human LC3. We examined the levels and distribution of a widely used Atg8a-GFP reporter transgene in retinal NDUFV1 knockdown animals and found increased reporter levels in retinas by quantitative immunofluorescence (Fig. 3A and B). We next examined autophagic flux to determine if increased levels of the Atg8a-GFP reporter were associated with dysfunctional autophagy. We assessed autophagic flux using a GFP-mCherry-Atg8a reporter in the brains of NDUFV1 with pan-neuronal knockdown mediated by the *elav-GAL4* driver and in control flies. The tandem fluorescent GFP-mCherry-Atg8a protein has been used to follow the maturation and progression of autophagosomes to autolysosomes in a variety of model systems (26,27), including *Drosophila* (28–30). When the GFP-mCherry-Atg8a protein is in the acidic autolysosome, GFP fluorescence is quenched, leaving only the mCherry signal. Thus, a small GFP to mCherry ratio indicates preserved autophagic flux, while a larger ratio indicates a flux deficit. The brains of NDUFV1 knockdown flies expressing the GFP-mCherry-Atg8a reporter displayed increased numbers of GFP-positive puncta that were also positive for mCherry (Supplementary Fig. 3A), as quantified by a higher GFP to mCherry ratio (Supplementary Fig. 3B), consistent with impaired autophagic flux. We then asked if complex I inhibition regulated the autophagy/lysosome system through a transcriptional mechanism. We performed quantitative real-time PCR for a panel of autophagy genes, including *Atg1, Atg4a, Atg5, Atg6, Atg7, Atg8a* and *Atg9*, in RNA prepared from heads of pan-neuronal NDUFV1 knockdown flies and controls. We were not able to confirm robust alterations of *Atg1, Atg5, Atg6, Atg7, Atg8a or Atg9*, but did find significantly elevated levels of *Atg4a* transcripts (Supplementary Fig. 3E).

**Figure 3.**
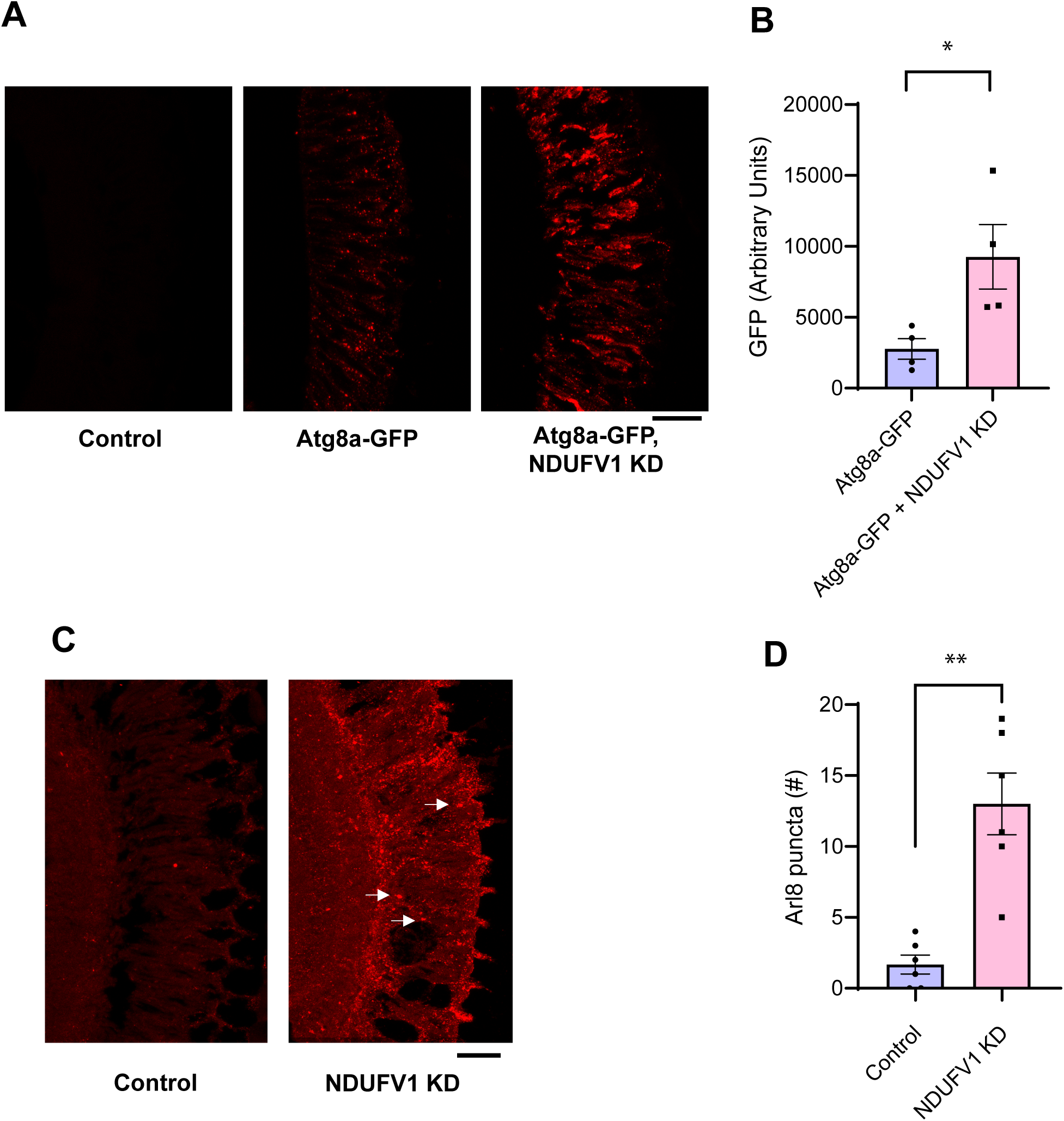
Increased autophagosome and lysosome markers following NDUFV1 knockdown. (A) Immunostaining for GFP in retinas from control flies and flies expressing the Atg8a-GFP reporter with and without knockdown of retinal NDUFV1 reveals increased reporter levels, consistent with altered autophagy. (B) Quantification of GFP immunofluorescence shows increased levels of the Atg8a-GFP reporter with retinal NDUFV1 knockdown. n = 4 per genotype. (C) Immunostaining for Arl8 in retinas from control and NDUFV1 knockdown retinas reveals lysosome staining with complex I inhibition, arrows. (D) Quantification shows an increase in Arl8 immunoreactive puncta in retinas with NDUFV1 knockdown compared to the controls. n = 6 per genotype. Data are represented as mean ± SEM. * p<0.01, ** p<0.002, t-test. Scale bars are 20 μm. Control is *GMR-GAL4/+.* Flies are 10 days old. Alt text: Photographs and bar graphs with quantification showing altered markers of the autophagosome/lysosome system following NDUFV1 knockdown.

Lysosomal changes frequently accompany dysfunctional autophagy. We first assessed lysosomes by immunostaining retinal complex I knockdown and control flies for Arl8, the *Drosophila* homolog of mammalian Arl8a and Arl8b, Arf-like GTPases localized to lysosomal membranes (31,32). Analysis of Arl8 staining revealed an increase in the number of lysosomes in complex I knockdown retinas compared to controls (Fig. 3C, arrows, and D). We additionally assessed the lysosomal compartment by labeling freshly dissected brains with LysoTracker Green. LysoTracker is a membrane-permeable dye, which selectively stains acidic compartments, including lysosomes. Increased numbers of LysoTracker-positive puncta were present in the brains of pan-neuronal NDUFV1 knockdown flies compared to controls (Supplementary Fig. 3C and D), consistent with an expanded lysosomal compartment in complex I deficient flies.

A major goal of developing a retinal toxicity model for complex I deficiency is to facilitate higher throughput approaches, including genetic screening, to investigate mechanisms underlying disease pathogenesis. We therefore performed a forward genetic screen to harness the power of forward genetics and identify mechanisms mediating proteostasis dysfunction in complex I deficiency. We focused on kinases because these enzymes are well conserved between *Drosophila* and mammals and represent attractive drug targets. Based on prior curation (33), we identified 358 kinases with available RNAi reagents for gene knockdown and crossed each RNAi line (Supplementary Table 1) to flies with knockdown of NDUFV1 in the retina using the *GMR-GAL4* driver (Fig. 4A). Retinal sections were prepared and assessed for histologic evidence of degeneration. We identified 13 preliminary suppressors and 13 preliminary enhancers in our screen. Since our goal is to ameliorate complex I toxicity we focused our subsequent efforts on the suppressors. The ability of a second, independent RNAi line to suppress complex I toxicity was tested to confirm the identity of suppressors. We thereby confirmed the identity of two kinases: pelle and RIOK1 (Fig. 4B). Of note, we independently identified RIOK2, a highly related kinase, as a suppressor. pelle is a serine/threonine protein kinase that functions in the Toll signaling pathway. RIOK1 is an atypical RIO kinase/ATPase with broad, but incompletely defined roles in cellular signaling (34).

**Figure 4.**
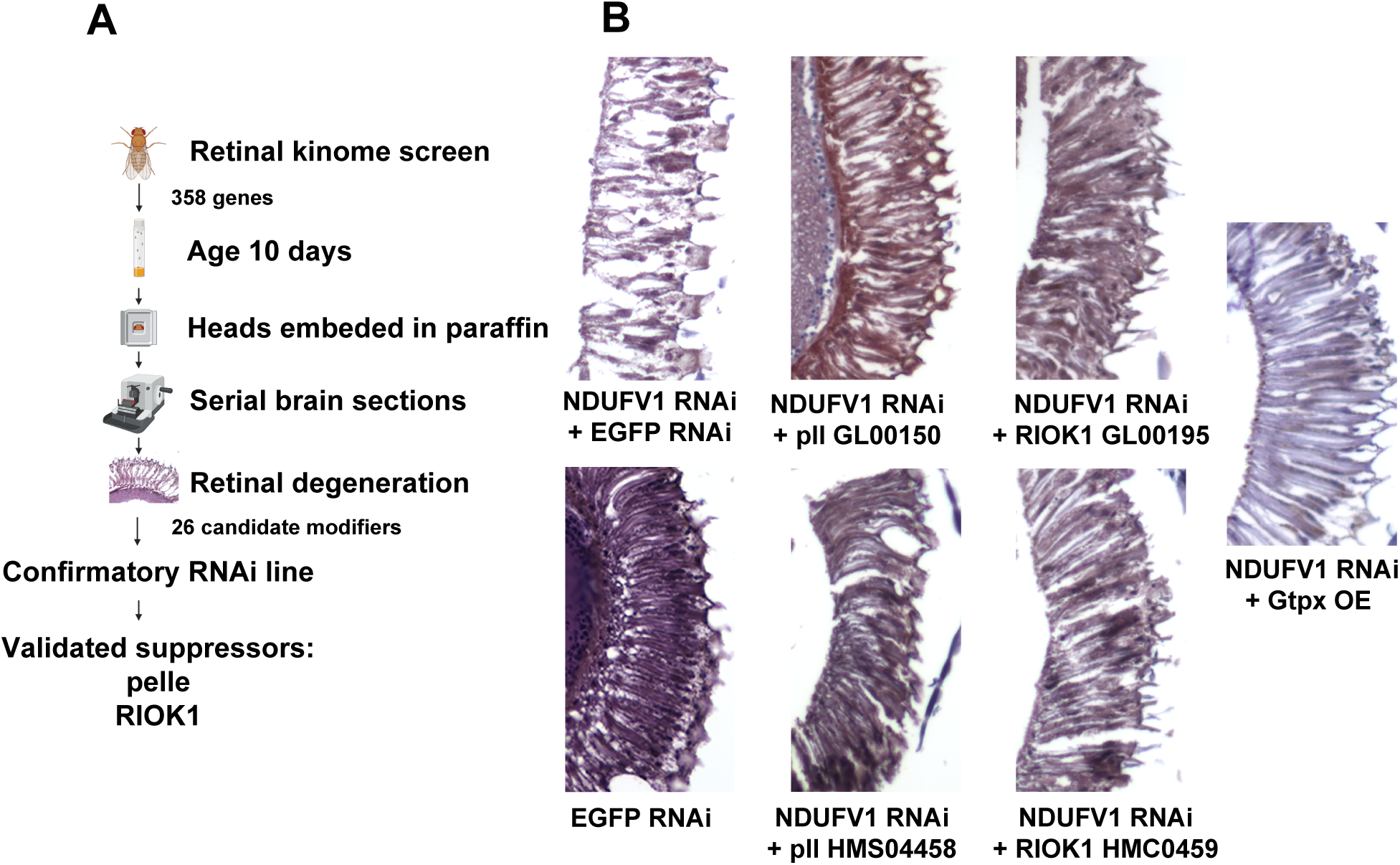
Rescue of complex I retinal toxicity with knockdown of pelle or RIOK1, or overexpression of Gtpx. (A) Schematic diagram of an in vivo transgenic RNAi whole kinome screen to identify genes and pathways mediating neurotoxicity of retinal complex I deficiency in aging neurons. The kinases pelle and RIOK1 were identified as loss of function suppressors of retinal degeneration caused by NDUFV1 knockdown. (B) Hematoxylin and eosin-stained sections of retinas from flies with retinal knockdown of NDUFV1 show degeneration, which is partially rescued by reduction of pelle or RIOK1 using two independent transgenic RNAi lines per gene, or by overexpression of the candidate oxidative stress modifier, glutathione peroxidase Gtpx. RNAi directed to *EGFP* is used as a control; driver is *GMR-GAL4.* Flies are 10 days old. Alt text: Schematic diagram of kinome screen and photographs showing rescue of retinal degeneration in NDUFV1 knockdown animals with genetic modifiers.

We were interested in expanding our genetic analysis of complex I-associated neurodegeneration and dysproteostasis beyond the modifiers identified in the kinome screen and reasoned that oxidative stress might play a role in complex I knockdown toxicity. The role of the electron transport chain in controlling levels of reactive oxygen species (4) is well documented. In addition, we identified several modifiers involved in the antioxidant defense system in the genome-scale pan-neuronal transgenic RNAi screen described above: GstD5 (Glutathione S transferase D5), Mgstl (Microsomal glutathione S-transferase-like) and Prx3 (peroxiredoxin 3). Consistent with these results we observed activation of a well-characterized oxidative stress reporter, GstD1-GFP (35), whose activation can be detected by immunostaining for GFP (Supplementary Fig. 4A and B) in retinal complex I knockdown flies. Overexpression of the glutathione peroxidase Gtpx is effective in alleviating oxidative stress (36) and rescuing neurotoxicity in fly models of human neurodegenerative diseases associated with mitochondrial dysfunction and increased oxidative free radicals (37–39). We thus increased levels of Gtpx to probe the functional role of oxidative stress in mediating cellular toxicity produced by complex I deficiency. We found that overexpression of Gtpx rescued NDUFV1 knockdown retinal toxicity (Fig. 4B).

We next determined if our genetic modifiers influenced NDUFV1 knockdown toxicity by altering proteostasis pathways. Retinal knockdown of pelle or RIOK1 did not significantly alter the effect of complex I-mediated inhibition on the levels of the unstable GFP-CL1 protein (Fig. 5A). In contrast, degradation of GFP-CL1 was increased in retinal NDUFV1 knockdown animal also overexpressing Gtpx (Fig. 5A). We then examined autophagosomes using an antibody recognizing endogenous Atg8a (28,40,41). As expected (Fig. 3), retinal complex I inhibition increased the number of Atg8a-immunoreactive puncta in the retina (Fig. 5B, arrows, and C) and pan-neuronal knockdown increased Atg8a-II by western blot analysis (Supplementary Fig. 4C and D). Knockdown of pelle in the retina did not significantly influence autophagosome number in retinal NDUFV1 knockdown animals (Fig. 5B, arrows, and C). RIOK1 knockdown in the retina reduced Atg8a staining, although autophagosome number did not return to control levels. In contrast, overexpressing Gtpx1 in the retina completely reversed the increased numbers of autophagosomes seen with complex I reduction (Fig. 5B and C). The optic lamina, which contains photoreceptor axons originating from retinal photoreceptors, was used for the Atg8a immunostaining experiments because the antibody had high background in the retina.

**Figure 5.**
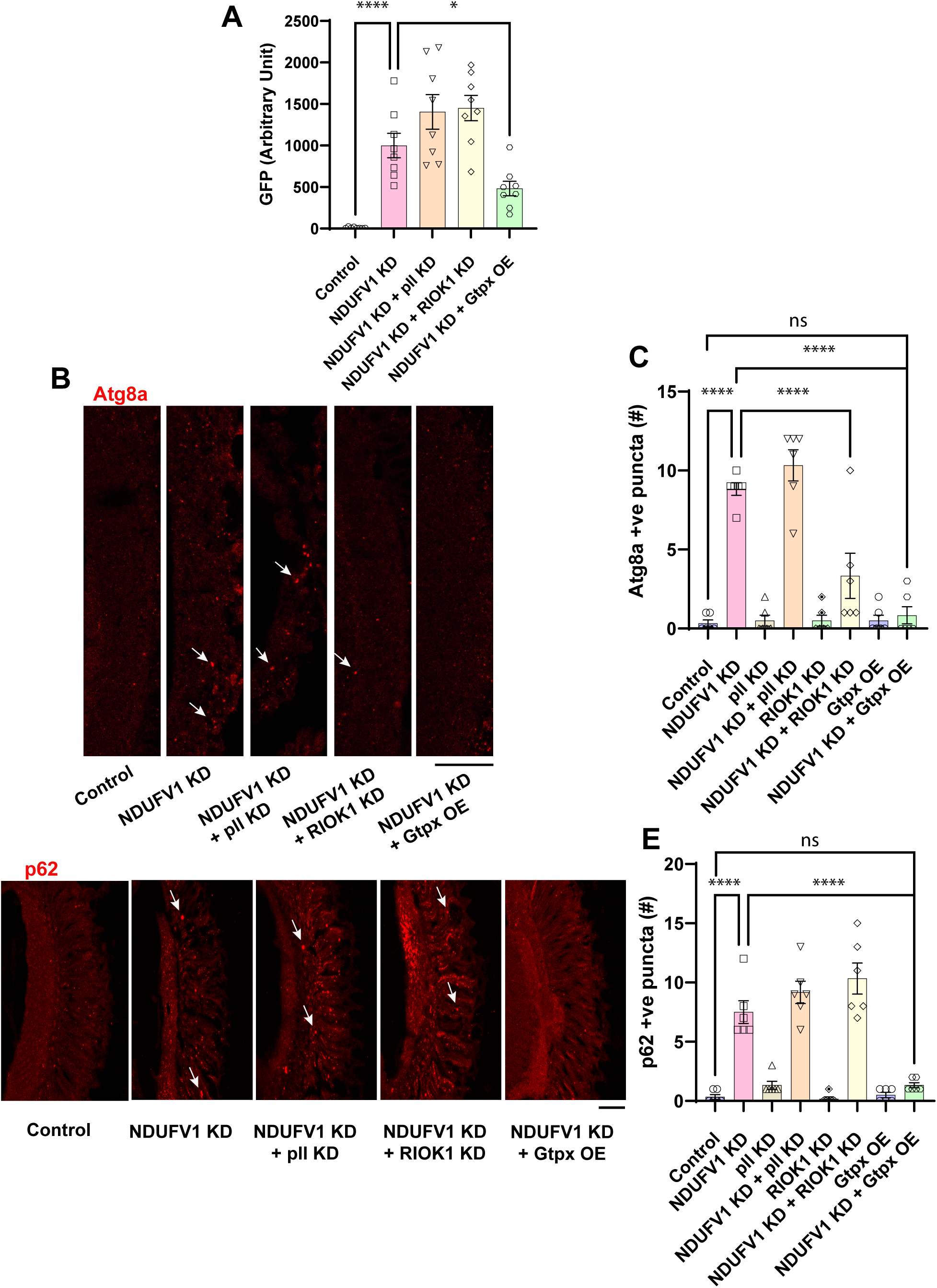
Effect of suppressors of NDUFV1 knockdown retinal toxicity on proteostasis pathways. (A) Quantification of GFP-CL1 western blots shows partial recovery of GFP degradation in retinal NDUFV1 knockdown flies with overexpression of Gtpx but no significant effect of pelle or RIOK1 RNAi. (B) Immunostaining for Atg8a (LC3) in the optic lamina from control flies and flies with retinal knockdown of NDUFV1 with and without RNAi directed to pelle or RIOK1 or overexpression of Gtpx shows normalization in the numbers of autophagosomes, arrows, by RIOK1 knockdown or Gtpx overexpression, but no influence of pelle knockdown. Atg8a-positive puncta are indicated by arrows. (C) Quantification of the number of Atg8a-positive puncta. (D) Immunostaining for p62 puncta, arrows, in retinas from control flies and flies with retinal knockdown of NDUFV1 with and without genetic modifiers shows reduction of puncta numbers in animals with complex I inhibition additionally overexpressing Gtpx, but no change with knockdown of pelle or RIOK1. (E) Quantification of the numbers of p62-positive puncta. Scale bars are 20 µm. n = 6 per genotype. Data are represented as mean ± SEM. *p<0.05, ****p<0.0001, ANOVA with Student-Newman-Keuls posthoc test. Control is *GMR-GAL4/+.* Flies are 10 days old. Alt text: photographs and bar graphs with quantification showing increased the effect of modifiers of NDUFV1 toxicity on proteostasis markers.

The p62 protein, also named sequestosome 1 (SQSTM1), binds to both ubiquitin and LC3 and acts as an adaptor to promote degradation of ubiquitinated proteins. Upon autophagic activation, p62 is recruited to autophagosomes and eventually degraded in lysosomes. p62 accumulation frequently accompanies aberrant autophagosome function and is commonly used as an indicator of autophagic flux (26,28). Immunostaining using an antibody directed to *Drosophila* p62, ref(2)P, revealed an increase in p62-positive puncta in the retina of flies with retinal NDUFV1 knockdown compared to controls, consistent with impaired autophagy (Fig. 5D, arrows, and E). Increased p62 was also seen in pan-neuronal NDUFV1 knockdown animals by western blotting (Supplementary Fig. 4E and F). Retinal knockdown of pelle or RIOK1 did not significantly alter the effect of complex I-mediated inhibition on the numbers of retinal p62-positive puncta (Fig. 5D, arrows, and E). In contrast, overexpressing Gtpx in the retina reduced the number of retinal p62 puncta back to control levels (Fig. 5D and E).

## Discussion

As part of ongoing studies of aging-dependent proteostasis failure and neurodegeneration we unexpectedly recovered components of the mitochondrial electron transport chain, including complex I and complex IV subunits. We found that knockdown of electron transport chain components produced age-dependent neurodegeneration and abnormal protein aggregation (Fig. 1). These findings raised the possibility that the pathogenesis of electron transport chain deficiency in patients, associated with a range of important human disorders from Leigh’s disease to retinal degeneration to myopathy, may include a defect in cellular proteostasis. To test this idea and facilitate a genetic approach to investigation of the mechanisms underlying the pathogenesis of electron transport chain deficiency, we created a flexible genetic model to study complex I deficiency by reducing the largest complex I component, NDUFV1, specifically in the aging adult retina.

When we reduced levels of NDUFV1 in the retina we observed progressive adult retinal degeneration (Fig. 4 and Supplementary Fig. 2). Retinal neurons are a key target of mitochondrial dysfunction (42,43). The large majority of inherited optic neuropathies are caused by mitochondrial dysfunction (44,45). The two most frequent inherited optic neuropathies are Leber’s hereditary optic neuropathy and dominant optic atrophy. Leber’s hereditary optic neuropathy is a respiratory chain deficiency disorder caused in most cases by mutations in the mitochondrial DNA-encoded complex I subunits ND1, ND4 and ND6. Heterozygous mutations in the gene encoding OPA1, a protein critical for normal mitochondrial fusion, underlie dominant optic atrophy. Mutations in genes encoding other mitochondrial quality control proteins, including mitofusin 2, DRP1 and paraplegin, cause syndromic optic neuropathy, further connecting mitochondrial function to retinal degeneration (44).

Metabolic vulnerabilities may underlie the sensitivity of retinal neurons to mitochondrial dysfunction. The retina is highly metabolically active (46). Proper electron transport chain function is not only critical to meet the substantial energy demand of the retina, but also to maintain redox balance and cytosolic pH, alterations of which can both lead to overproduction of reactive oxygen species (47). Indeed, oxidative stress is a consistent feature of cellular and animal models of complex I deficiency based on Leber’s hereditary optic neuropathy mitochondrial mutations, in some cases without clear reduction in ATP levels (48). Given the strong implication of oxidative stress in the pathogenesis of complex I deficiency we assessed the ability of increasing expression of the glutathione peroxidase Gtpx to influence neurotoxicity in our retinal model of complex I deficiency. Glutathione peroxidases form a key component of endogenous antioxidant systems by reducing H_2_O_2_ and organic hydroperoxides to water and corresponding alcohols (36,49). Increased levels of Gtpx protect from oxidative stress in vivo produced by paraquat feeding (36) or by expression of toxic human tau in neurodegenerative tauopathy models (38). Here we show that Gtpx overexpression is highly effective at suppressing retinal degeneration associated with NDUFV1 knockdown (Fig. 4), consistent with a role for oxidative stress mediating complex I loss of function neurotoxicity. These findings are further consistent with recovery of the closest human homolog of Gtpx, GPX4, as a synthetic lethal in a genome-wide CRISPR screen for modulators of mitochondrial chemical inhibitors (50).

Interestingly, rescue of retinal degeneration by Gtpx overexpression was correlated with normalization of proteasome and autophagy markers (Fig. 5). Thus, ameliorating oxidative stress may promote normalization of proteostasis by coordinately regulating two major arms of the proteostasis network. Formation of oxidative adducts that promote protein misfolding and aggregation provides the most direct connection between oxidative stress and dysproteostasis. Oxidative damage to key chaperones may provide may be especially damaging by potentiating the deleterious effects of oxidative adducts on the function protein folding machinery (51). In addition, the accumulation of oxidatively damaged proteins and other macromolecules can activate cellular stress signaling pathways that control levels and function of the ubiquitin/proteasome and autophagy/lysosomal systems. For example, the transcription factor Nrf2 plays a key role in the response to oxidative stress by regulating the expression of a set of antioxidant and detoxification enzymes and also transcriptionally activates proteasome subunit genes as well as multiple autophagy pathway genes (52). We investigated potential transcriptional regulation of autophagy in our complex I deficiency model by performing quantitative real time PCR for a panel of autophagy genes. Our approach failed to identify robust changes in most of the genes we studied but we did observe a two-fold increase in Atg4a transcript levels in NDUFV1 knockdown animals compared to controls (Supplementary Fig. 4E). The Atg4 family of proteases plays a key role in autophagy by cleaving and activating Atg8 proteins (53). Altered regulation of Atg4 may thus play a role in autophagic dysfunction following complex I inhibition. Overall, further work will be needed to characterize the mechanisms responsible for proteostasis control by oxidative stress in the current complex I deficiency model.

Our findings connecting mitochondrial dysfunction with oxidative stress and altered proteostasis in retinal degeneration secondary to complex I deficiency fit well within the broader context of ocular disease pathogenesis. Aberrant proteostasis has been strongly implicated in the age-related disorders macular degeneration, cataract and glaucoma (54). Alterations in the ubiquitin/proteasome and autophagy/lysosome systems have been documented in various genetic forms of retinal degeneration and in other aging-dependent ocular diseases (54,55), as has oxidative stress and mitochondrial dysfunction (42,43), suggesting fundamental connections among these pathways. A functionally important relationship between altered proteostasis and photoreceptor vulnerability was recently suggested by a chemical screening approach in an induced pluripotent stem cell-derived model of retinal degeneration caused by genetic ciliary dysfunction, which identified reserpine as both promoting photoreceptor survival and normalizing proteostasis (56).

Genetic complex I deficiency syndromes include not only disorders, like Leber’s hereditary optic neuropathy, with a dominant clinical syndrome of retinal degeneration, but also encompass a broad array of other clinical presentations as well. Encephalopathy, termed Leigh’s disease, and myopathy are common primary manifestations of genetic complex I deficiency. The retina is also affected in many of these patients as well. Patients with complex I deficiency caused by nuclear gene defects frequently have ocular abnormalities and optic atrophy is one of the most specific signs pointing to complex I deficiency in affected children. Unfortunately, prognoses are poor for affected individuals. About half of patients diagnosed with complex I deficiency die before the age of two years, with only a quarter of patients reaching 10 years (57). Current treatments are primarily supportive with no clear disease-modifying yet available (6). There is thus an urgent need for new approaches to identifying clinically targetable mechanisms.

Given the clinical relevance of retinal degeneration in complex I deficiency patients we have developed here a retinal model of complex I deficiency in the powerful genetic model organism *Drosophila*. From a technical perspective, restricting complex I knockdown to the retina, a nonessential tissue in the fly, allows us to use a variety of genetic reagents to knockdown gene expression that would be lethal if employed in a ubiquitous or pan-neuronal fashion. Although the *Drosophila* system facilitates many useful approaches to studying gene function and identifying therapies, including candidate testing (Fig. 4B) and drug screening (17), one of the most powerful applications is forward genetic screening. Here we have capitalized on the forward genetic screening approach in two ways. First, we identified altered proteostasis as a candidate mechanism mediating complex I deficiency nervous system toxicity in a genome-scale screen (Fig. 1). Second, we performed a targeted kinome screen in which we knocked down the complement of protein kinases (Supplementary Table 1) and identified RIO kinases and pelle as loss of function suppressors retinal degeneration in complex I deficiency (Fig. 4).

The RIO (*<u>ri</u>*ght *<u>o</u>*pen reading frame) family of proteins are classified as atypical kinases because they have broad structural homology to the ancestral family of eukaryotic protein kinases, but lack overall sequence similarity (58). There are two RIO kinases in unicellular organisms, and an additional RIOK family member in multicellular eukaryotes (59). RIOK1 and RIOK2 have conserved roles in ribosomal RNA biogenesis and cell cycle progression (34). Additional roles for RIO kinases have been suggested in diverse pathways, including protein arginine methylation (60), regulation of p53 activity and stability (61,62), and AKT (61) and p38 MAPK (63,64) signaling.

In the roundworm *C. elegans* RIOK1 acts upstream of the p38 MAPK signaling pathway to negatively regulate innate immunity (64). Interestingly, the other kinase recovered in our screen, pelle, has a key role in the innate immune system. The NF-kB signaling pathway is conserved from *Drosophila* to mammals and plays a central role in innate immunity in invertebrates and vertebrates. In mammals, proinflammatory cytokines of the IL-1 family bind to Toll-like receptors, which signal through a receptor complex containing the IL-1R-associated kinase (IRAK). Signal activation promotes autophosphorylation and dissociation of IRAK from the receptor complex leading through a kinase cascade to nuclear translocation and activation of NF-kB (65). The *Drosophila* ortholog of IRAK is pelle. Thus, the RIO kinases and pelle may modulate complex I neurotoxicity through innate immune signaling. Alternatively, another known, or previously undescribed, pathway regulated by the kinases may be critical for mediating the effect on retinal degeneration in our model.

Our findings that, unlike Gtpx overexpression, knockdown of either RIOK1 or pelle, does not consistently normalize markers of the ubiquitin/proteasome and autophagy/lysosomal systems in NDUFV1 knockdown animals (Fig. 5) suggests that the effects of RIOK1 and pelle reflect the activity of different pathways than those controlled by Gtpx. In particular, the inability of RIOK1 or pelle knockdown to influence the stability of the GFP-CL1 protein suggests that neither kinase protects from neurotoxicity arising in complex I deficiency by increasing proteosome activity. In contrast, increased expression of Gtpx, which plausibly acts by reducing oxidative stress, appears to enhance both the activity of both the ubiquitin/proteasome and autophagy/lysosome systems. Although further work will be required to define the molecular pathways underlying the protective effects of RIOK1 and pelle knockdown, our findings raise the possibility that coordinate therapeutic targeting of the oxidative stress and immune signaling pathways may be beneficial in complex I deficiency patients. Despite extensive evidence in preclinical models and patient tissues for a role of oxidative stress in diverse neurologic disorders, including complex I deficiency, antioxidants have been disappointing in the clinic. As in cardiovascular disease (66), combination therapies targeting multiple pathways underlying disease pathogenesis may be an attractive treatment approach.

In addition to suggesting the potential utility for combination therapies, our findings may have other implications for therapy development. We chose to perform a kinome screen because these enzymes are traditionally amenable to small molecule modulation. Indeed, inhibitors of RIO kinases are under development (67,68). The role of RIO kinases in ribosome function raises the concern regarding potential toxicity of RIOK inhibition, although the ability of RIOK knockdown to rescue retinal degeneration in vivo (Fig. 4) suggests that therapeutic levels of inhibition might be achieved without significant toxicity. Finally, our results may have implications beyond genetic complex I deficiency syndromes. Mitochondrial dysfunction has been implicated in the pathogenesis of a broad range of neurologic disorders, including age-related neurodegenerative diseases. Loss of normal protein homeostasis is a hallmark of aging (69) and protein aggregates are an invariant feature of common human neurodegenerative disorders, including Parkinson’s disease, Alzheimer’s disease and related disorders. Thus, the mechanisms and therapeutic strategies suggested here in the context of complex I deficiency may have broader applicability.

## Materials and Methods

### Drosophila stocks

*Drosophila* crosses were performed at 25°C. All flies were aged at 25°C and were assayed at 1, 10 or 30 days as noted in the text and figure legends. Both male and female flies were used in each experiment. Transgenic RNAi lines for the genome-scale and kinome screens and GAL4 driver lines were obtained from the Bloomington *Drosophila* Stock Center. The genome scale screen was performed on flies aged for 30 days, as will be described in detail elsewhere (Leventhal, Fraenkel, Feany, in preparation). The kinome screen was performed on 10-day-old flies. The following additional stocks were also obtained from the Bloomington *Drosophila* Stock Center: *UAS-COX10/CG5073 JF01522*, *UAS-COX10 JF01671*, *UAS-NDUFV1/ND-51 HM05213, UAS-NDUFV1/ND-51 HMJ21591, UAS-NDUFV1/ND-51 HMS01590, UAS-Dcr2; UAS-EGFP shRNA, UAS-GFP-mCherry-Atg8a*. Other stocks were kind gifts of the indicated investigators: *UAS-Gtpx*, F. Missirlis; *UAS-Atg8a-GFP*, Thomas Neufeld; *GstD1-GFP*, Dirk Bohmann; *UAS-GFP-CL1* and *UAS-GFP-G76V*, Udai Pandey.

#### Histology, Immunostaining and Imaging

*Drosophila* heads were fixed in formalin and embedded in paraffin before sectioning. Serial frontal sections (2 or 4 μm) of the entire brain were prepared and mounted on glass slides. To assess brain and retina morphology, sections were stained with hematoxylin and eosin. For immunostaining on paraffin sections antigen retrieval was performed by microwaving in sodium citrate buffer for 15 minutes. Slides were then blocked in 2% milk in PBS with 0.3% Triton X-100, followed by overnight incubation with primary antibodies at room temperature. Primary antibodies to the following proteins were used at the indicated concentrations: GABARAP/Atg8a (1:1,000, E1J4E, Cell Signaling), Arl8 (1:1000, Developmental Studies Hybridoma Bank), GFP (1:50 to 1:2,000, N86/8, Neuromab), p62/ref(2)P (1:1,000, our laboratory). After three washes with PBS-Triton, slides were incubated with appropriate secondary antibodies coupled to Alexa Fluor 488, Alexa Fluor 555 or Alexa Fluor 647. Rabbit polyclonal antibodies directed against *Drosophila* p62 (ref(2)P) were created by Covance using the peptide sequence PRTEDPVTTPRSTQ (30). Quantification of Atg8a and p62 in Figure 5 was performed by immunostaining with biotinylated secondary antibodies, followed by avidin–biotin–peroxidase complex (Vectastain Elite; Vector Laboratories), as previously described (23). GFP reporters were detected using immunostaining for GFP because endogenous fluorescence was inadequate for detection and quantification.

Imaging of fluorescent markers was performed with Zeiss laser-scanning confocal microscopy. Transgenic flies expressing GFP-mCherry-Atg8a were examined to assess autophagic flux. Brains from 10-day-old animals were dissected in PBS and imaged immediately. For quantification a region of interest was selected and the intensity of GFP and mCherry were measured using ImageJ as described (29,30). LysoTracker staining was performed following incubation of freshly dissected brains of 10-day-old flies in 0.05 μM LysoTracker for 5 minutes at room temperature. Brains were mounted in PBS and imaged immediately. The number of puncta per 1500 μm^2^ was counted. For other markers a region of interest was defined and total fluorescence or the number of puncta were determined. All images were taken using 63X-oil lens with 1.3X zoom and processed in Zen Blue software. The laser intensity was the same across all genotypes during acquisition and each channel was tested for saturation across all stacks. A sequential scan was performed to avoid any crosstalk between fluorophores. Post-acquisition, the intensity of all the images were modified in an identical manner across genotypes.

#### Molecular biology

RNA was isolated from 10-day-old fly heads using QIAzol (QIAGEN). Following RNA isolation, complementary DNA was prepared using the Applied Biosystems High-Capacity cDNA Reverse Transcription Kit using the manufacturer’s protocol. SYBR Green (Applied Biosystems) based qPCR was performed on an Applied Biosystems QuantStudio 6 Flex Real-Time PCR System. Primer sequences for Atg4a were GCG CTC TTC GAG ATC AGT CA (forward) and CCT GCC GCT CTC TTC AAC TA (reverse). Primer sequences for RpL32, used as internal control, were GAC CAT CCG CCC AGC ATA C (forward) and CGG CGA CGC ACT CTG TT (reverse). The fold change in gene expression was determined by the ΔΔCt method, where Ct is the threshold value.

#### Immunoblotting

For western blot analysis *Drosophila* heads were homogenized in 15 μl of 2x Laemmli buffer (Sigma-Aldrich). Samples were boiled for ten minutes and subjected to SDS-PAGE. Protein were transferred onto nitrocellulose membranes (Bio-Rad), blocked in 2% milk in PBS with 0.05% Tween 20 (PBSTw) and incubated overnight at 4°C with primary antibodies. Membranes were washed three times with PBSTw and incubated for three hours with appropriate HRP-conjugated secondary antibody (SouthernBiotech) at room temperature. Following multiple washes with PBSTw, proteins were visualized by enhanced chemiluminescence (Alpha Innotech). Protein transfer and loading were monitored by Ponceau S staining. Blots were reprobed with an antibody directed to actin to illustrate equivalent protein loading in figures. All immunoblots were repeated at least three times. Primary antibodies to the following proteins were used at the indicated concentrations: actin (1:1,000, JLA20, Developmental Studies Hybridoma Bank), GFP (1:1,000, JL-8, Clontech), GABARAP/Atg8a (1:2,000, E1J4E, Cell Signaling), Ref2P/p62 (1:3,000, 178440, Abcam).

## Abbreviations

GFP: green fluorescent protein;
RIOK: right open reading frame kinase;
family NDUFV1: NADH:ubiquinone oxidoreductase core subunit V1;
ATP: adenosine triphosphate;
GMR: glass multimer reporter;
GstD5: Glutathione S transferase D5;
Mgstl: Microsomal glutathione S-transferase-like;
Prx3: peroxiredoxin 3.

## Acknowledgements

Fly stocks were obtained from the Bloomington *Drosophila* Stock Center (NIH P40-OD018537), and Drs. F. Missirlis, T. Neufeld, D. Bohmann, and U. Pandey. We thank the Transgenic RNAi Project (TRiP) at Harvard Medical School (NIH R24 OD030002; PI: N. Perrimon) for making *Drosophila* stocks (70). Monoclonal antibodies were obtained from the Developmental Studies Hybridoma Bank developed under the auspices of the NICHD and maintained by the University of Iowa, Department of Biology, Iowa City, IA 52242, and the UC Davis/NIH NeuroMab Facility. Yi Zhong provided excellent technical assistance. This research was funded by the Department of Defense (PR160270), and in whole or in part by Aligning Science Across Parkinson’s ASAP-000301 through the Michael J. Fox Foundation for Parkinson’s Research (MJFF).

---

**Supplementary Figure 1.**
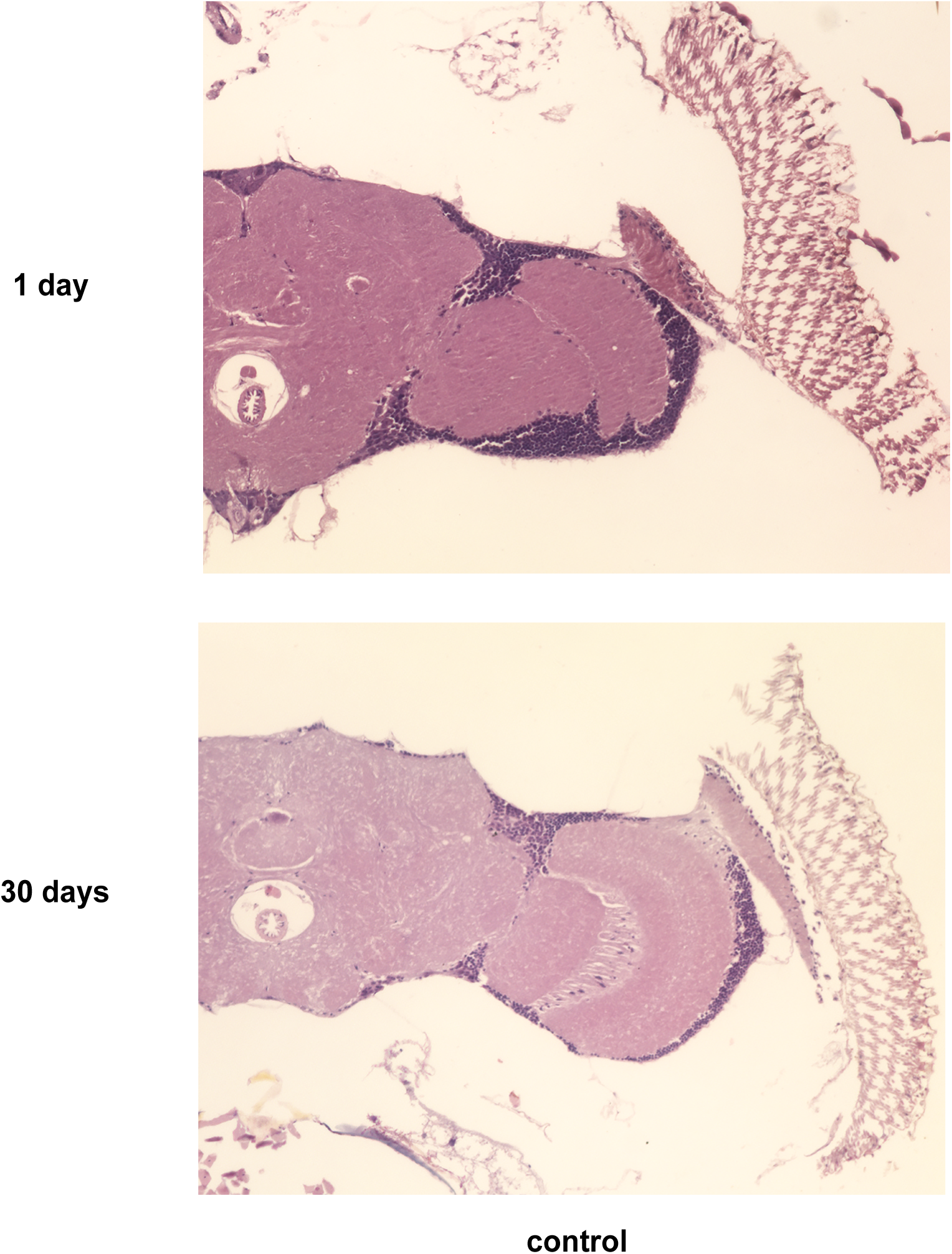
Control brains are well-preserved at 30 days of age. Hematoxylin and eosin-stained sections of brains from control flies at 1 and 30 days of age demonstrate overall maintenance of brain and retinal structure with age. Genotype (control) is *elav-GAL4/+.* Flies are the indicated ages.

**Supplementary Figure 2.**
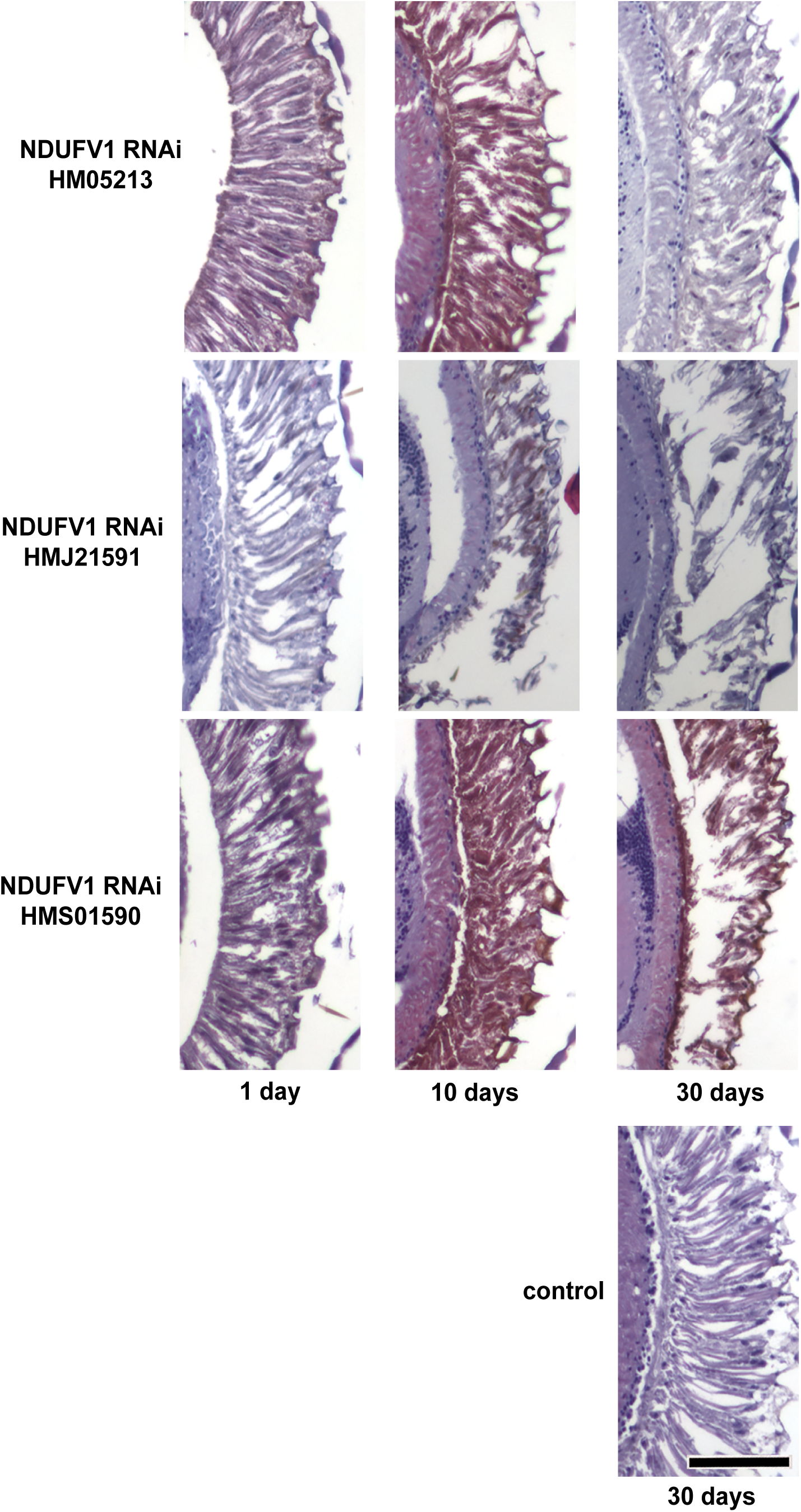
Retinal degeneration with knockdown of NDUFV1 with age. Hematoxylin and eosin-stained sections of retinas from flies with retinal knockdown of NDUFV1 by three independent RNAi lines showing retinal degeneration with age. Scale bar is 50 µm. Control is *GMR-GAL4/+.* Flies are the indicated ages.

**Supplementary Figure 3.**
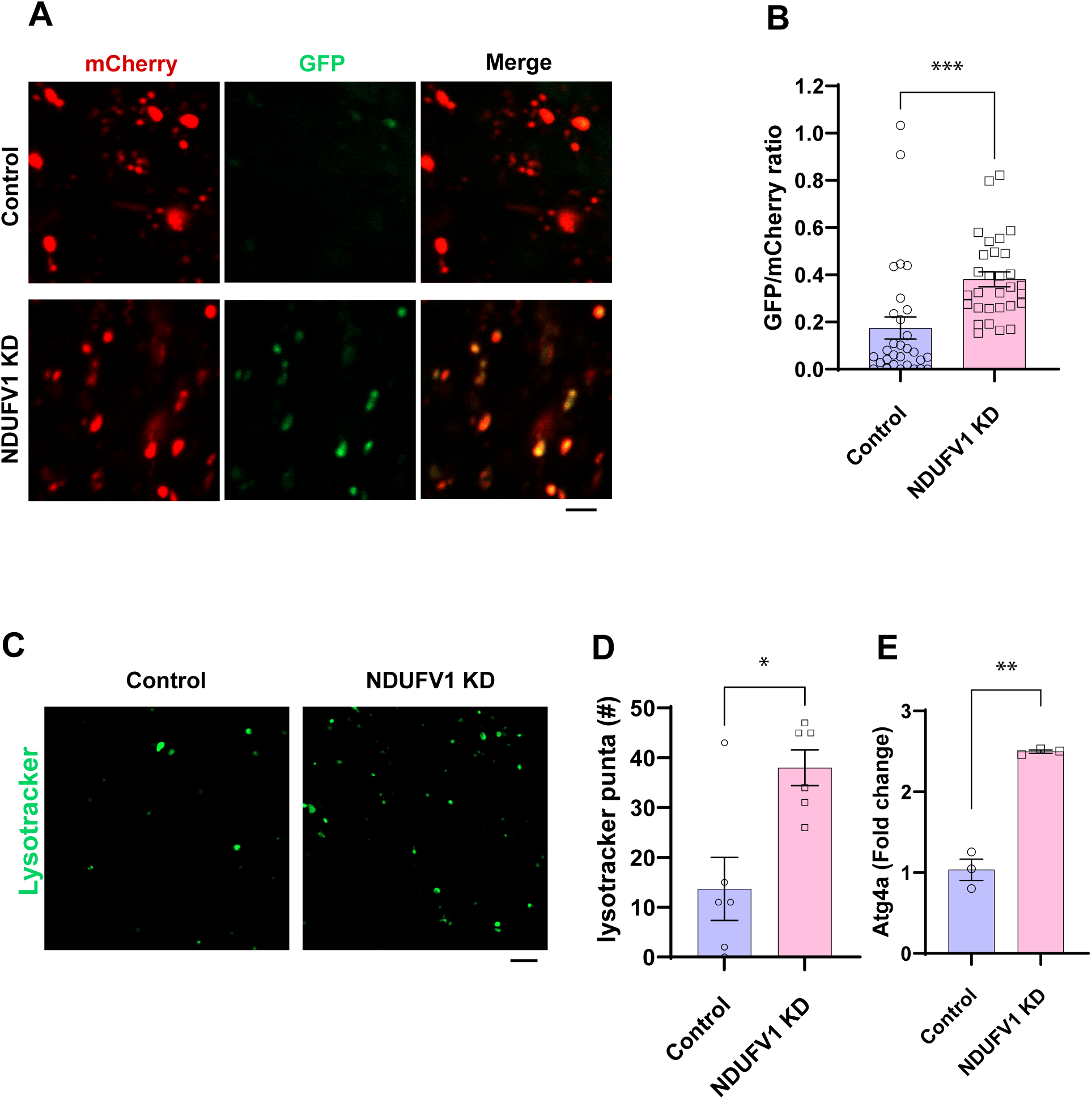
Autophagic and lysosomal changes in the brains of flies with pan-neuronal NDUFV1 knockdown. (A) Representative images from flies expressing the tandem reporter GFP-mCherry-Atg8a with and without pan-neuronal knockdown of NDUFV1. (B) Quantification shows an increase in the GFP to mCherry fluorescence ratio in the brains of pan-neuronal NDUFV1 RNAi-expressing flies, indicating impaired autophagic flux. n = 30 puncta from 3 animals per genotype. Control is *elav-GAL4/+; UAS-GFP-mCherry-Atg8a/+* in A and B. (C) Images show lysosomes stained with LysoTracker. (D) Quantification shows an increase in stained lysosomes in the brains with pan-neuronal NDUFV1 knockdown. n = 6. (E) Real-time PCR shows an increase in *Atg4a* transcript levels with pan-neuronal NDUFV1 knockdown. n = 3 repeats with 10 flies per repeat. Control is *elav-GAL4/+* in C, D and E. Data are represented as mean ± SEM. *p<0.05, **p<0.01, ***p < 0.001, t-test. Scale bars are 2 μm in A and 5 μm in C. Flies are 10 days old.

**Supplementary Figure 4.**
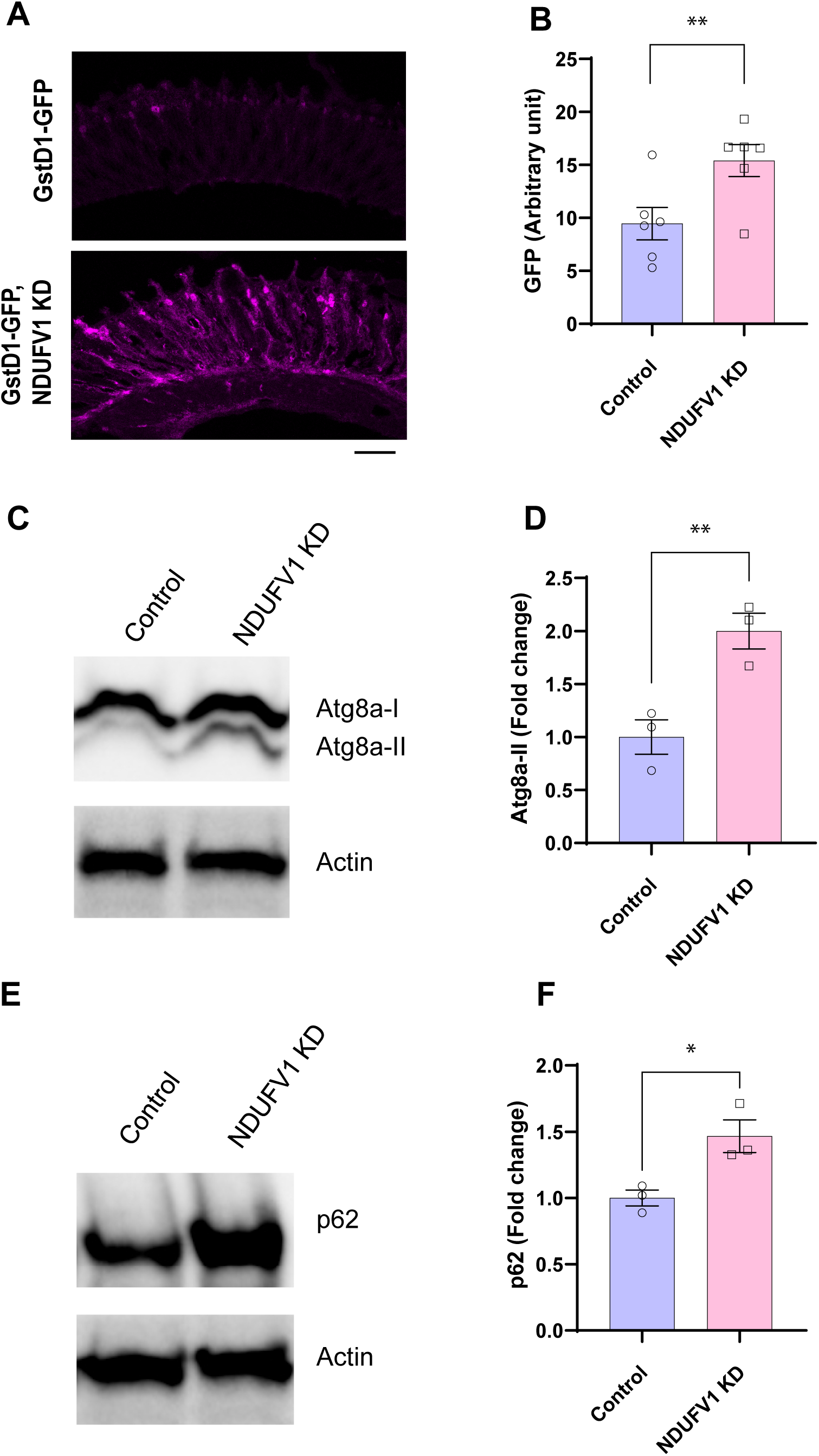
Increased oxidative stress with retinal NDUFV1 knockdown. (A) Immunostaining for GFP in retinas from flies expressing *GstD1-GFP*, an oxidative stress reporter, in the retina with and without retinal NDUFV1 knockdown. Scale bar is 20 μm. (B) Quantification shows the increase in oxidative stress reporter levels due to NDUFV1 knockdown. n = 6 per genotype. Control is *GMR-GAL4/+; GstD1-GFP/+*. Flies are 10 days old. (C – F) Western blot (C, E) and quantification (D, F) shows increased Atg8a-II (C, D) and p62 (E, F) in pan-neuronal NDUFV1 knockdown fly heads. n = 3 per genotype. Control is *elav-GAL4/+.* Flies are 30 days old. All data are represented as mean ± SEM. *p<0.05, **p<0.01, ***p < 0.001, t-test.

